# Pro-coagulant lipids in physiological ratios found in the activated platelet membrane do not impact clot structure or fibrinolysis in purified assays

**DOI:** 10.1101/2025.08.11.669632

**Authors:** Bethan H Morgan, Laura E Farleigh Smith, Daniela O Costa, Victoria J Tyrrell, Josefin Ahnström, Peter Vincent Jenkins, Peter W Collins, Nicola J Mutch, Valerie B O’Donnell

## Abstract

**Purpose:** A central role for the pro-coagulant membrane comprising aminophospholipids (aPL) and enzymatically oxidized phospholipids (eoxPL) in promoting hemostasis via interaction with coagulation factor Gla domains is well established. However, little is known about their interactions with the fibrinolytic pathway, their ability to alter clot structure or to support the activated protein C (APC) pathway. Previous studies used membrane liposome compositions that differ from those expected physiologically and/or generated inconsistent findings. To address this, pro-coagulant membranes comprising physiological proportions of aPL and eoxPL will be tested for their ability to support fibrinolysis using standard assays.

**Methods:** The impact of phospholipids on clot structure and clot lysis was tested using absorbance-based assays. To investigate the impact of PS or eoxPL on fibrinolysis, plasmin was monitored chromogenically, and clot dissolution measured in a purified lysis system activated by tissue plasminogen activator or urokinase. To determine the impact of eoxPL on APC/protein S, FVa was incubated with APC (+/−protein S) in a purified prothrombinase assay.

**Results:** At the concentrations of lipids tested in our study, PS did not significantly impact clot structure or fibrinolysis. Similarly, eoxPL did not impact either fibrinolysis or activity of APC/Protein S.

**Conclusion:** Using liposome compositions that approximate activated blood cells, we found that the pro-coagulant membrane is unlikely to influence either clot structure or fibrinolytic activity directly, beyond its well characterized role in supporting Gla dependent coagulation factors and the actions of platelet associated proteins/receptors.

## Introduction

Phospholipids (PL) are key constituents of plasma membranes and play essential roles in coagulation.^1,2^ In resting platelets, the aminophospholipids (aPL), comprising phosphatidylserine (PS) and phosphatidylethanolamine (PE), are primarily located on the inner membrane, but upon platelet activation they are externalized to the outer side.^1,3–5^ Coagulation involves the generation of thrombin (FIIa), and follows the sequential activation of a series of enzymes and co-factors (Figure 1).^6,7^ Several of these enzyme/co-factor complexes feature γ-carboxyglutamic acid-rich (Gla) domains that interact with aPLs and calcium, both of which are essential for their function.^2,8^ Here, Gla domains bind to PS and calcium, while PE acts to support this.^9^ As well as the Gla domain containing coagulation factors, FV and FVIII are also known to interact with phospholipid membranes via their C domains.^2^

**Figure 1.**
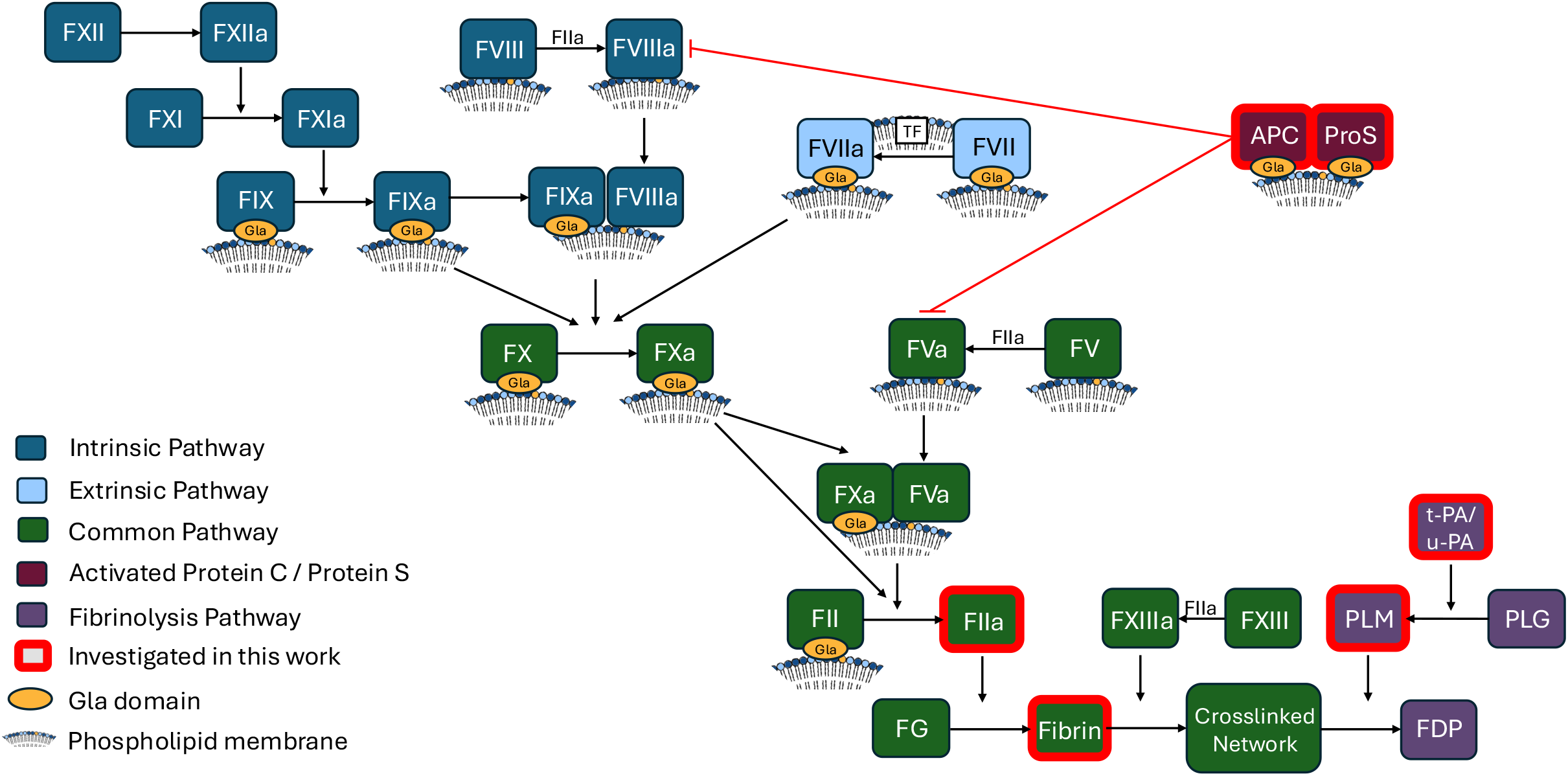
Overview of the coagulation cascade including fibrinolysis and activated protein C pathways, highlighting phospholipid interactions. TF, Tissue factor; APC, activated protein C; ProS, protein S; FG, fibrinogen; PLM, plasmin; PLG, plasminogen; t-PA, tissue plasminogen activator; u-PA, urokinase; FDP, fibrin degradation products.

In addition to the head groups, fatty acyl (FA) chains of PL can also regulate thrombin generation. In particular, enzymatically oxidized PL (eoxPL), which contain an oxygenated FA at the Sn2 position can enhance coagulation.^10,11^ For these lipids, it has been proposed that steric interference from the electronegative oxygenated FA increases space between headgroups, increasing the accessibility of PS and calcium ions for Gla domains.^12,13^ While the external membrane of resting platelets is mainly phosphatidylcholine (PC), after platelet activation, external aPL varies from 4 – 30%, depending on the activating stimulus, while it has been estimated that < 5-10 % of PE are eoxPL.^1,13^

Several natural anticoagulants also contain Gla domains, and as a result their activities can be influenced by membrane lipid composition. Activated protein C (APC) is a Gla domain containing enzyme which exerts an anticoagulant function by inactivating FVa and FVIIIa. PS has previously been demonstrated to support APC function, additionally requiring PE for optimal activity.^14^ APC activity is greatly enhanced by its Gla domain containing co-factor protein S. Protein S has a higher binding affinity for lipid membranes than APC, and forms a complex with APC and FVa on the membrane, which enhances APC activity.^15^ Furthermore, preliminary studies previously showed that oxidized lipids derived from brain tissue can enhance APC activity.^16^ However, whether physiological levels of eoxPL from platelets and leukocytes can enhance APC, with/without protein S, is unknown.

Aside from interaction with Gla domain containing factors, whether the pro-coagulant membrane could also play a regulatory role in downstream pathways, influencing fibrin structure and clot lysis are less clear, with published studies being contradictory.^17–24^ Thrombin cleaves fibrinogen to fibrin which rapidly forms a network whose structure dictates function. A fibrin clot formed of thin fibers is mechanically stiff and can impede blood flow or fracture leading to thrombotic events, while a clot comprised of fewer fibers, interspersed by large pores, may be too weak to seal the wound.^25–27^ During network formation, fibrin monomers aggregate laterally to form protofibrils.

These protofibrils then undergo further lateral aggregation to form the basis of the fibers within the network. The subsequent increase in fiber diameter can be measured by absorbance.^28–30^ Using this approach, studies using synthetic membranes suggested that PL can influence clot structure by interacting with fibrinogen, however, the membrane compositions tested were not representative of the platelet surface.^22–24^ One of these studies additionally suggested that thrombin could directly interact with di-palmitoyl-PC (DPPC), regulating its activity,^23^ however, DPPC is not an abundant platelet PC, and furthermore the PS binding Gla domain of prothrombin is lost during activation to thrombin. Thus, it’s not clear how aPL could directly impact thrombin enzymatic activity.^31^

Fibrinolysis describes the process of clot dissolution after hemostasis is complete. Here, tissue plasminogen activator (t-PA) and urokinase (u-PA) convert plasminogen to its active form plasmin, which then cleaves fibrin causing breakdown of the fiber network.^32,33^ A few studies have investigated whether pro-coagulant aPL directly impact the fibrinolytic system, but these have yielded conflicting results.^17–21^ The conditions used to measure fibrinolysis vary substantially, and the lipid compositions tested are generally non-physiological. For example, studies on t-PA activation of fibrinolysis used membranes comprising up to 100% PS.^20^ OxPL have also been found bound to plasminogen, leading to suggestions they may play a role in fibrinolysis.^34^ However, whether they can impact fibrinolysis when incorporated into the platelet membrane is currently unknown.

Clot structure, fibrinolysis and anti-coagulation are essential for the regulation of hemostasis, and their dysregulation is a direct contributor to human pathology, thus it is important to clarify whether known procoagulant lipids have a direct impact on these parameters.^35,36^ To address this, a series of approaches will be used, focusing on lipid compositions that are representative of physiological membranes. First, the impact of the procoagulant membrane on thrombin activity will be tested to exclude any impact of lipids on the subsequent bioassays, which require consistent thrombin activity. Then the impact of liposomes containing procoagulant lipids on clot structure will be investigated. Next, the impact of liposomes containing physiological proportions of PC, PE, PS, and eoxPL, to represent the activated platelet surface, will be investigated in plasmin activity assays and the effect of these membranes on fibrinolysis will be measured in a purified system. Last, whether eoxPL generated by immune cells can regulate inactivation of FVa by APC in both the presence and absence of protein S will be tested. In this study, the impact of aPL and eoxPL will be tested using purified systems in place of plasma. This is because in complete plasma the impact of coagulation itself would not be possible to control for, introducing a confounding factor. Specifically, thrombin would be generated at variable amounts (regulated by lipids), and could independently influence clot structure, fibrinolysis and FVa generation.^7,26^ Thus, our experimental approach using purified proteins specifically allows us to directly test for an effect of aPL and eoxPL on clot structure, plasmin activity and activation, and the inhibition of FVa by APC with and without protein S without interference from endogenous thrombin generated in plasma.

## Materials and methods

### Generation and preparation of lipids

In order to test the effect of aPL and eoxPL on clot structure, fibrinolysis, and APC activity liposomes containing varying amounts of the aPL 1-stearoyl-2-oleoyl-sn-glycero-3-phospho-L-serine (SOPS) and two forms of eoxPL, 15-hydroxyeicosatetraenoic acid-PE (15-HETE-PE) and 15-hydroxyeicosatetraenoic acid-PC (15-HETE-PC) were used as described in Table 1. 15-HETE-PE and 15-HETE-PC were generated using soybean 15-lipoxygenase (15-LOX) (Santa Cruz Biotechnology, Dallas, Texas, USA) as described previously.^37^ Briefly 1-stearoyl-2-arachidonoyl-sn-glycero-3-phosphoethanolamine (SAPE) (Avanti Polar Lipids, Alabaster, Alabama, USA) or 1-stearoyl-2-arachidonoyl-sn-glycero-3-phosphocholine (SAPC) (Avanti Polar Lipids, Alabaster, Alabama, USA) were incubated with 15-LOX in deoxycholate buffer in the presence of O_2_. Lipid hydroperoxides were then reduced using SnCl_2_ and the lipids extracted. 15-HETE-PE or 15-HETE-PC were separated from the residual SAPE or SAPC using HPLC and the purity confirmed using mass spectrometry. The concentrations of 15-HETE-PE and 15-HETE-PC were calculated from their absorbance at 235 nm using the extinction coefficient 28 mM^-1^cm^-1^.

**Table 1.** Liposome compositions used in experimental work.

| Label | Lipid Composition |
| --- | --- |
| Buffer | - |
| 0% PS | 50% DSPC, 30% SOPE, 20% SOPC |
| 5% PS | 50% DSPC, 30% SOPE, 15% SOPC, 5% SOPS |
| 10% PS | 50% DSPC, 30% SOPE, 10% SOPC, 10% SOPS |
| 20% PS | 50% DSPC, 30% SOPE, 20% SOPS |
| PC/PE | 65% DSPC, 20% SOPE, 10% SAPE, 5% SOPC |
| PC/PE/PS<sup>a</sup> | 65% DSPC, 20% SOPE, 10% SAPE, 5% SOPS |
| PC/PE/PS(15-HETE-PE)<sup>a</sup> | 65% DSPC, 20% SOPE, 10% 15-HETE-PE, 5% SOPS |
| PC/PE/PS<sup>b</sup> | 55% DSPC, 10% SAPE, 20% SOPE, 10% SAPC, 5% SOPS |
| PC/PE/PS(15-HETE-PE)<sup>b</sup> | 55% DSPC, 10% 15-HETE-PE, 20% SOPE, 10% SAPC, 5% SOPS |
| PC/PE/PS(15-HETE-PC) | 55% DSPC, 10% SAPE, 20% SOPE, 10% 15-HETE-PC, 5% SOPS |

**Notes:** <sup>a</sup> used in all experiments apart from APC/protein S, <sup>b</sup> used for APC/protein S experiments.
**Abbreviations:** DSPC, 1,2-distearoyl-sn-glycero-3-phosphocholine; SOPE, 1-stearoyl-2-oleoyl-sn-glycero-3-phosphoethanolamine; SOPC, 1-stearoyl-2-oleoyl-sn-glycero-3-phosphocholine; SOPS, 1-stearoyl-2-oleoyl-sn-glycero-3-phospho-L-serine; SAPE, 1-stearoyl-2-arachidonoyl-sn-glycero-3-phosphoethanolamine; 15-HETE-PE, 15-hydroxyeicosatetraenoic acid-PE; SAPC, 1-stearoyl-2-arachidonoyl-sn-glycero-3-phosphocholine; 15-HETE-PC, 15-hydroxyeicosatetraenoic acid-PC.

Lipids were added to a glass vial at the ratios given for each condition in Table 1 and dried under N_2_ gas. The lipids used were as follows: 1,2-distearoyl-sn-glycero-3-phosphocholine (DSPC), 1-stearoyl-2-oleoyl-sn-glycero-3-phosphocholine (SOPC), 1-stearoyl-2-oleoyl-sn-glycero-3-phosphoethanolamine (SOPE), 1-stearoyl-2-arachidonoyl-sn-glycero-3-phosphoethanolamine (SAPE), 1-stearoyl-2-arachidonoyl-sn-glycero-3-phosphocholine (SAPC) and SOPS (Avanti Polar Lipids, Alabaster, Alabama, USA), and 15-HETE-PE and 15-HETE-PC as generated above.

Lipids were re-suspended in 500 μl liposome buffer (20 mM HEPES and 100 mM NaCl, pH 7.35) to give a total lipid concentration of 100 μM, then vortexed and subjected to ten freeze thaw cycles, and passed through a 100 nm pore membrane 19 times using a LiposoFast extruder (Avestin, Inc. Ottawa, Canada). All liposome preparations were used on the day of generation.

The lipids selected for these experiments were chosen with the aim of creating a system suitable for testing the effect of PS and eoxPL, which would be expected to provide a simple representation of the surface of an activated platelet. The liposome composition was varied around a standard ratio of 65% PC, 30% PE and 5% PS which reflects a standard cell membrane surface.^1,13^ DSPC was used as it represents an abundant form of PC found in mammalian cell membranes. SAPE and SAPC were used to act as a control for 15-HETE-PE and 15-HETE-PC which feature an oxidized FA chain derived from the arachidonic acid in SAPE and SAPC. SOPE, SOPC, and SOPS were used as they are not susceptible to oxidation and would therefore not be expected to interfere with the evaluation of eoxPL. For measurements of thrombin activity and clot structure the composition included 50% DSPC, 30% SOPE and up to 20% SOPC substituted with SOPS. This allowed for alteration of the headgroup without influencing the FA chains. For fibrinolysis experiments the liposomes contained 65% DSPC, 20% SOPE, 10% SAPE and 5% SOPC where the SOPC was again substituted with SOPS, and SAPE was substituted with 15-HETE-PE. These compositions tested the substitution of PC with PS without influencing the FA composition, and substitution of the arachidonic acid from SAPE with 15-HETE without otherwise modifying the liposomes. Similarly for APC/protein S experiments SAPC and SAPE were substituted with 15-HETE-PC and 15-HETE-PE respectively.

### Thrombin activity assay

To assess thrombin activity, final concentrations of 0.56 mM of Chromogenix S-2238 (Enzyme Research Laboratories, Swansea, UK) in Tris buffered saline with tween (TBST, 10 mM Tris, 140 mM NaCl and 0.01% Tween-20 at pH 7.4) and 60 μM liposomes were combined in a 96 well plate. 5 mM CaCl_2_ and 0.25 NIH/ml human alpha thrombin (Enzyme Research Laboratories, Swansea, UK) in TBST were added immediately before loading into a Clario Star Plus plate reader (BMG Labtech, Aylesbury, UK) to a total volume of 100 μl. After a 2 s double orbital shake at 500 rpm, absorbance (405 nm) was monitored for 80 min at 37°C. The gradient of absorbance during the linear period (0-20 min) was calculated to give the Δ Absorbance (405 nm) per minute.

### Clot structure assay

To measure clot structure, final concentrations of 2 mg/ml plasminogen-depleted fibrinogen (Enzyme Research Laboratories, Swansea, UK) in TBST and 60 μM liposomes were combined in a 96 well plate. 5 mM CaCl_2_ and 0.25 NIH/ml human alpha thrombin in TBST were added to the wells immediately before loading into a Clario Star Plus plate reader, to a total volume of 100 μl. After a 2 s double orbital shake at 500 rpm, absorbance at 405 nm was measured every 30 s at 37°C until a plateau was reached. The maximum absorbance (405 nm) during the plateau region following clot formation was recorded.

### Plasmin activity assay

To measure the effect of liposomes on plasmin activity, 50 μl liposomes of varying concentrations (0-20 μM) were added to a 96 well plate along with a final concentration of 12.5 nM plasmin (Enzyme Research Laboratories, Swansea, UK). 500 μM Chromogenix S-2251 (Quadratech Diagnostics, Eastbourne, UK) in TBST was then added to the wells, immediately before loading into a Clario Star Plus plate reader, to a total volume of 200μl. After a 2 second double orbital shake at 500 rpm, absorbance at 405 nm was measured every minute for 6 h at 37°C. The gradient of the absorbance data during the linear period (0-100 min) was calculated to give the Δ Absorbance (405 nm) per minute.

### Fibrinolysis assay

To determine the effect of liposomes on fibrinolytic activity, liposomes of varying concentrations (0-20 μM) were added to a 96 well plate (20 μl). Final concentrations of 1.25 mg/ml fibrinogen, 0.24 μM human glu-plasminogen (Enzyme Research Laboratories, Swansea, UK) and either 600 pM native human urokinase (u-PA) (Abcam, Cambridge, UK) or 20 pM Hyphen BioMed recombinant human t-PA (Quadratech Diagnostics, Eastbourne, UK) in TBST were added to each well. 5 mM CaCl_2_ and 0.25 NIH/ml thrombin were then added immediately before loading into a Clario Star Plus plate reader, to a total volume of 100μl. After a 2 s double orbital shake at 500 rpm the absorbance (405 nm) was measured every minute for 6 h at 37°C. Lysis time is defined as the time between 50 % of maximum absorbance during formation to 50 % maximum absorbance (405 nm) during breakdown of the network, derived by interpolation between the points of the raw data traces.

### FVa inactivation assay

Phospholipids (20 μM) were incubated at 37°C in the presence of FVa (5 nM) (Prolytix, Essex Junction, Vermont, USA), CaCl_2_ (5 mM) (Merck, Dorset, UK), with/without protein S (50 nM) (Prolytix, Essex Junction, Vermont, USA) and with/without APC (0.25 nM) (Enzyme Research Laboratories, Swansea, UK) in a total volume of 30 μL. The assay was performed in buffer containing 20 mM HEPES (Sigma Aldrich, Dorset, UK) 0.1 M sodium chloride (Sigma Aldrich, Dorset, UK) and 0.01 % BSA (Sigma Aldrich, Dorset, UK) at pH 7.4. After 5 minutes, APC was inactivated by addition of 4-Amidinophenylmethanesulfonyl fluoride hydrochloride (p-APMSF) (0.2 mM) (Santa Cruz Biotechnology, Dallas, Texas, USA)^38^ and incubated at room temperature for 20 min. FVa activity remaining was measured using a prothrombinase assay. For this, the remaining samples from the FVa inactivation assays were topped up with liposomes to a total final concentration of 25 μM using the control liposomes (PC/PE/PS^b^). Prothrombin (500 nM) (Enzyme Research Laboratories, Swansea, UK) and FXa (20 nM) (Enzyme Research Laboratories, Swansea, UK) were then added to a final total volume of 50 μL and samples incubated at 37°C for 5 min. NOTE: The concentration of total FVa is diluted to 3 nM and the concentration of CaCl_2_ is diluted to 3 mM for this step. The reaction was quenched by addition of 12.5 μl of 35 mM EDTA (final concentration 7 mM) (Sigma Aldrich, Dorset, UK). The concentration of thrombin generated was measured by comparing the initial linear rate of cleavage of the Chromogenix substrate S-2238 (Enzyme Research Laboratories, Swansea, UK) by the samples to the rate of cleavage using a set of thrombin standards of known concentrations (FIIa) (Enzyme Research Laboratories, Swansea, UK). Absorbance was measured using a Clario Star Plus plate reader. Following a 10 s double orbital shake at 500 rpm, absorbance (405 nm) was measured every 38 s for 20 min at room temperature.

### Statistical analysis

For each condition three independent liposome preparations (n = 3) were analyzed, with each of these measured in triplicate (technical replicates). Statistical significance was determined using one-way ANOVA and Tukey’s post hoc test. Data processing and analysis was performed using Microsoft Excel and MATLAB.

## Results

### PS containing liposomes do not significantly influence thrombin activity

Prior to conducting assays testing the impact of PL on clot structure or lysis, it was important to ensure that under our conditions, lipids were not impacting thrombin activity. Although the formation of thrombin is PS dependent, the Gla domain of prothrombin is lost when prothrombin fragment 1.2 is released during activation to thrombin.^31^ While it was expected that PS containing liposomes would not impact thrombin activity, a previous study suggested that the activity could be enhanced by DPPC.^23^ To exclude this potential confounder, the ability of thrombin to cleave a chromogenic substrate in the presence of liposomes was evaluated. The liposomes tested approximated a physiological membrane, containing 70% PC (50% DSPC, 20% SOPC), and 30% PE as SOPE, to which gradually increasing amounts of SOPS up to 20% were incorporated, by replacing the SOPC.^1^ Replacing SOPC with SOPS ensured that the FA composition was consistent across all conditions. Thrombin activity was not significantly enhanced by the inclusion of lipids, and furthermore there was no impact of PS at any level tested (Figure 2 A,B; Supplementary Table 1).

**Figure 2.**
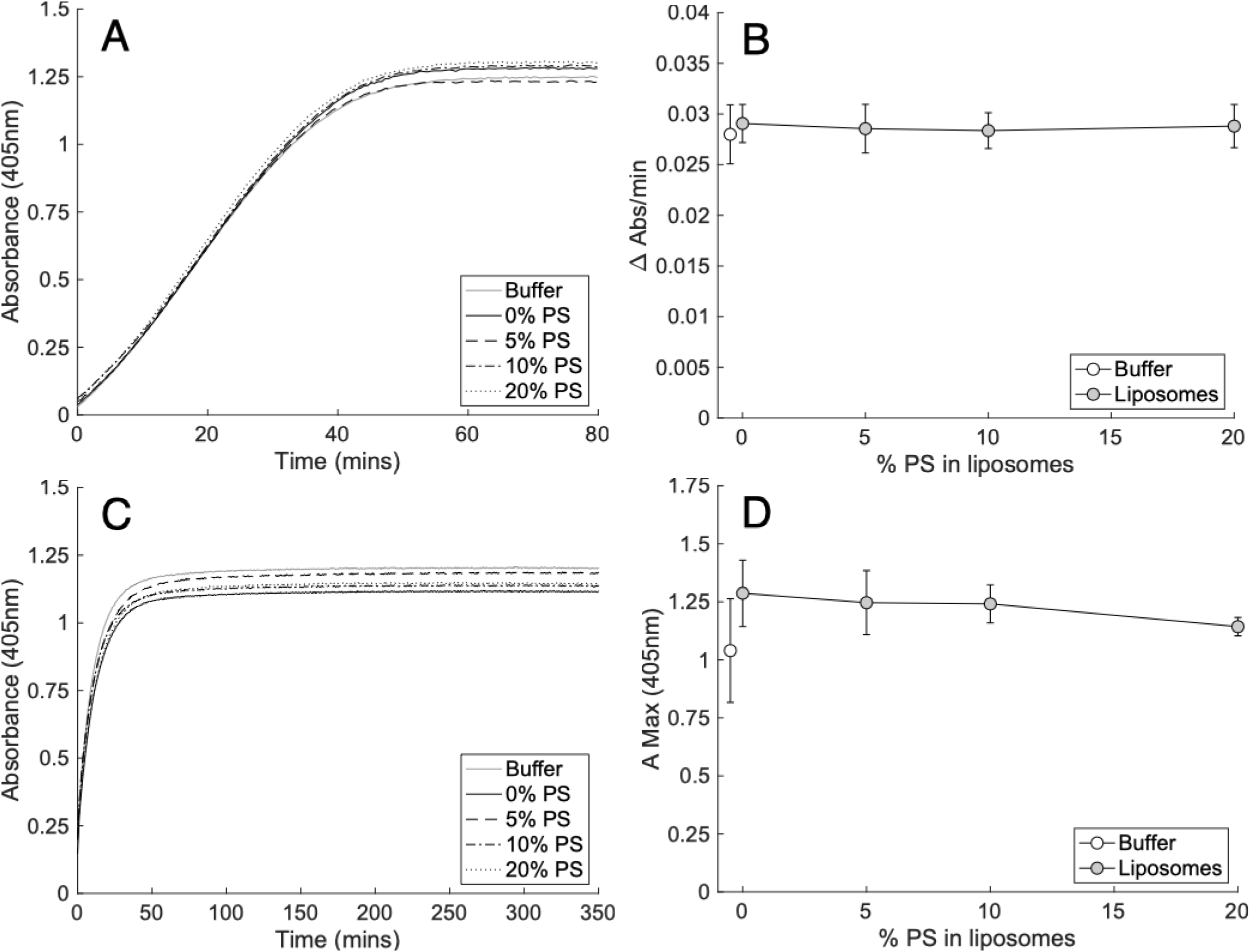
Liposomes containing physiological amounts of PS do not alter thrombin activity or structure of clots formed by purified thrombin cleavage of fibrinogen. Panels (A-B). PS did not impact the activity of 0.25 NIH/ml (A,B) thrombin. Thrombin activity was measured by the metabolism of its chromogenic substrate S-2238 (0.56 mM) in the presence of liposomes (60 μM) containing varying proportions of PS as outlined in methods (n = 3, mean ± SD). Panels (C-D). PS did not impact the structure of clots formed from fibrinogen (2 mg/ml) and 0.25 NIH/ml (C,D) thrombin. Clot structure was measured by the maximum absorbance (405 nm) of clots formed in the presence of liposomes (60 μM) containing a varying proportion of PS as outlined in methods (n = 3, mean ± SD). Left panels; representative traces, right panels; summary data. Statistics performed using one-way ANOVA. Composition of liposomes was as follows: 0% PS (50% DSPC, 30% SOPE, 20% SOPC), 5% PS (50% DSPC, 30% SOPE, 15% SOPC, 5% SOPS), 10% PS (50% DSPC, 30% SOPE, 10% SOPC, 10% SOPS) 20% PS (50% DSPC, 30% SOPE, 20% SOPS).

### PS did not significantly influence clot structure

The effect of membranes containing biologically-relevant ratios of PL, with varying PS amounts up to 20%, were next tested for their impact on the developing clot structure, measured by maximum absorbance at 405 nm.^28,29^ As shown, the clot structure was not impacted by PE/PC liposomes, either without PS or with gradually increasing amounts up to 20 % (Figure 2 C,D; Supplementary Table 2).

### Pro-coagulant lipids do not regulate plasmin activity

To test the effect of the pro-coagulant membrane on fibrinolysis, the activity of plasmin was first measured using a chromogenic assay in the absence of fibrin, while incorporating liposomes of varying composition. As above, liposomes were formulated to provide physiological ratios of PC/PE/PS, ensuring that the FA composition remained constant as PLs were substituted. We previously demonstrated that eoxPL are pro-coagulant, using ~65-70 % PC, 30 % PE, and 5– 10% PS liposomes, with eoxPL added up to 10 % maximum to model platelet membranes.^13^ Similarly, for this experiment, 5% SOPC (control) was replaced with SOPS to test the impact of PS. Also, to test the impact of eoxPL, 10 % SAPE (control) was replaced with its oxygenated analogue, the eoxPL, 1-stearoyl-2-15-HETE-PE (15-HETE-PE).^11,39^ Liposome concentrations were varied from 0 – 20 μM, to alter the total surface area tested. PE/PC membranes with or without PS had no impact on plasmin activity. Similarly, inclusion of HETE-PE had no effect at any concentration tested (Figure 3 A,B; Supplementary Table 3).

**Figure 3.**
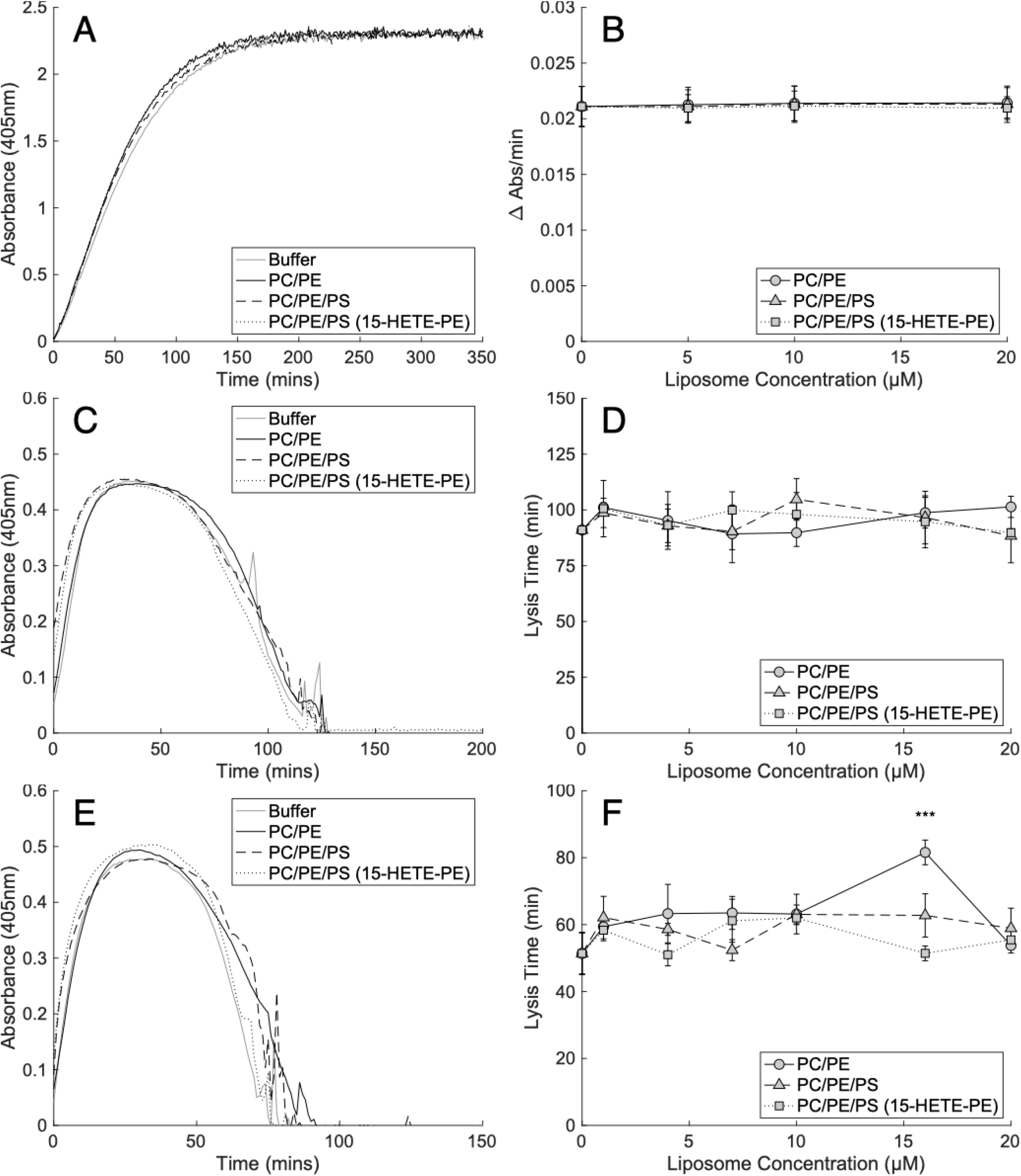
Pro-coagulant lipids have little or no impact on the activity of plasmin or its formation from plasminogen by u-PA or t-PA. Panels (A,B). Pro-coagulant liposomes did not impact the activity of plasmin. Plasmin (12.5 nM) activity was measured by the metabolism of its chromogenic substrate S-2251 (500 μM) in the presence of varying concentrations of pro-coagulant liposomes (0-20 μM) as outlined in methods (n = 3, mean ± SD). Panels (C,D) Liposomes did not alter lysis time of purified fibrin clots by plasminogen activated by t-PA. Lysis time was measured as the time from half maximal absorbance (405 nm) during clot formation to half maximal absorbance during the lysis of fibrin clots (1.25 mg/ml fibrinogen) formed by thrombin (0.25 NIH/ml) in the presence of CaCl_2_ (5 mM) where plasminogen (0.25 μM) was activated by t-PA (20 pM) as outlined in methods (n = 3, mean ± SD). Panel (C) shows liposomes at 4 μM. Panels (E,F) Liposomes did not alter lysis time of purified fibrin clots by plasminogen activated by u-PA, apart from at one single concentration. Lysis time was measured as the time from half maximal absorbance (405 nm) during clot formation to half maximal absorbance during the lysis of fibrin clots (1.25 mg/ml fibrinogen) formed by thrombin (0.25 NIH/ml) in the presence of CaCl_2_ (5 mM) where plasminogen (0.25 μM) was activated by u-PA (600 pM) as outlined in methods (n = 3, mean ± SD). Panel (E) shows liposomes at 10 μM. Statistics were performed using one-way ANOVA with Tukeys post-hoc test (* p<0.05, ** p<0.01, *** p<0.001). Composition of liposomes was as follows: PC/PE (65% DSPC, 20% SOPE, 10% SAPE, 5% SOPC), PC/PE/PS (65% DSPC, 20% SOPE, 10% SAPE, 5% SOPS), PC/PE/PS(15-HETE-PE) (65% DSPC, 20% SOPE, 10% 15-HETE-PE, 5% SOPS).

### Pro-coagulant lipids do not alter fibrinolysis

We next investigated whether aPL and eoxPL could regulate upstream activation of plasminogen and downstream dissolution of fibrin. Here, t-PA or u-PA were added to activate plasminogen in a fibrinolysis assay where liposomes were present along with fibrinogen, which was then clotted by the addition of thrombin. Formation/lysis of clots was determined using absorbance at 405 nm, as a measure of the presence of fibrin networks.^28,29^ In this assay, the formation and subsequent lysis is visualized by the shape of the absorbance curve, as shown in Figure 3 C. When t-PA was used as the plasminogen activator, liposomes comprising PE/PC had no impact at concentrations up to 20 μM. Furthermore, inclusion of PS with/without HETE-PE at physiological amounts had no impact (Figure 3 C,D; Supplementary Table 4). Similarly, when lysis was initiated by u-PA there was little effect on formation or lysis of the clot from the addition of pro-coagulant membranes (Figure 3 E,F; Supplementary Table 5). While a statistically significant prolongation of lysis time was observed for one condition, it occurred in the absence of PS, and only at a single liposome concentration (Figure 3 E,F; Supplementary Table 5).

### EoxPL phospholipids do not alter FVa inactivation by APC

It has previously been reported that air oxidized brain phospholipids can enhance APC activity.^16^ However, the molecular species in the mixture that were responsible for this were not determined. Considering this, we next tested whether eoxPL could regulate APC/protein S, similarly to how they regulate the activities of procoagulant factors, focusing on 15-HETE-PE and 15-HETE-PC.^12,13^ As above, a small amount of PE or PC from the control condition that contained arachidonic acid at Sn2 position was removed and replaced with the equivalent 15-HETE-PE or PC, using amounts reflective of platelet membrane compositions. There was no significant effect of liposomes containing either 15-HETE-PE or 15-HETE-PC on either FVa inactivation by APC alone, or on the enhancement of APC activity by protein S (Figure 4; Supplementary Table 6).

**Figure 4.**
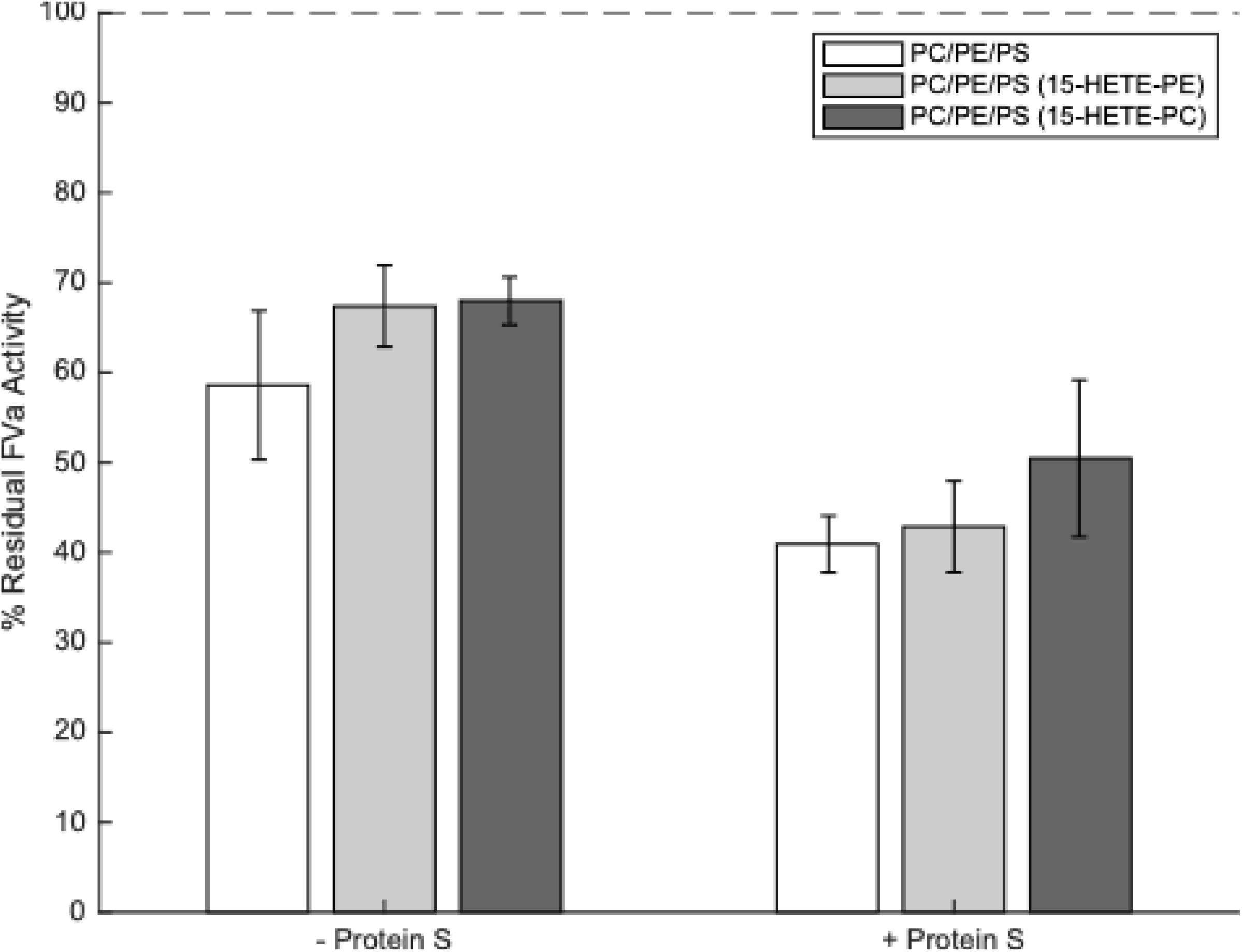
Liposomes containing 15-HETE-PE and 15-HETE-PC do not enhance activated protein C (APC) or protein S activities. Liposomes (20 μM, containing eoxPL as listed in Table 1), FVa (5 mM), CaCl_2_, (5 mM), APC (0.25 nM) +/−protein S (50 nM) were incubated at 37°C for 5 minutes, APC then inactivated with p-APMSF before remaining FVa activity measured using a prothrombinase assay (n = 3, mean ± SD). Results are recorded as a percentage of remaining FVa activity compared with controls containing no APC or protein S (I.e. 100 % FVa activity). Blanks that included no FVa were also tested to account for the level of FXa activity in the absence of FVa and these values were deducted from all other values. No significant differences between liposome types were detected, either in the presence or absence of protein S (tested by one-way ANOVA). Composition of liposomes was as follows: PC/PE/PS (55% DSPC, 10% SAPE, 20% SOPE, 10% SAPC, 5% SOPS), PC/PE/PS(15-HETE-PE) (55% DSPC, 10% 15-HETE-PE, 20% SOPE, 10% SAPC, 5% SOPS), PC/PE/PS(15-HETE-PC) (55% DSPC, 10% SAPE, 20% SOPE, 10% 15-HETE-PC, 5% SOPS).

## Discussion

The roles of PS and eoxPL in driving coagulation via their interactions with factors containing Gla domains is well established (Figure 1). However, whether aPL or eoxPL play a role in regulating clot structure or lysis is less clear. The existing literature on effects of the pro-coagulant lipid membrane on these elements of hemostasis is limited, utilize non-physiological liposome compositions, and in the case of fibrinolysis, are contradictory, while a role for eoxPL has not yet been investigated.^17–24^ Therefore, here we focused on determining the effect of physiologically relevant liposome compositions on fibrin clot structure and plasmin activation and activity.

Additionally, we investigated the effect of eoxPL on the Gla domain containing APC and protein S, which act as part of the inhibitory system limiting coagulation, and are already known to interact with PS.^2^

To investigate the effect of aPL on fibrin network structure and the breakdown of the fibrin network by plasmin, purified protein assays were used. This was to ensure that there were no confounding effects from changes in thrombin generation and activity during the assay. As expected, we confirmed experimentally that there was no effect of PS on thrombin activity under our assay conditions.

In previously published studies, synthetic PL membranes composed of PC and phosphatidylglycerol (PG) were reported to impact clot structure, as measured by absorbance-based assays, suggesting PL can directly interact with fibrinogen. ^22–24^ The mechanism was proposed to involve the adsorption of fibrinogen onto liposome surfaces. However, the composition of the lipid membrane was non-physiological, with inclusion of up to 20% PG, which is a low abundance lipid in platelets. Fibrinogen was also suggested to adsorb to PC/PE and PC/PS membranes.^22–24^ In these studies, the greatest effects were seen using non-physiological liposomes that contained a single PL molecular species, e.g. 100% dipalmitoylphosphatidylcholine (DPPC) or 100% 1-palmitoyl-2-oleoyl-sn-glycero-3-phosphatidylserine (POPS). Importantly, membranes containing physiologically-relevant ratios of both PE and PS, which directly model the pro-coagulant membrane, were not tested.^22–24^ Our study directly addressed this by using physiologically relevant lipid membranes comprising PC/PE and varying amounts of PS (0-20%) and found that these liposomes had no effect on mature clot structure. Overall, our data indicate that pro-coagulant membranes that form during platelet activation are unlikely to directly influence mature fibrin clot structure. However, in vivo, it is possible that lipids could indirectly contribute, through regulation of thrombin generation during the process of coagulation itself (Figure 1).

Clot dissolution is mediated by plasmin, which is generated in the blood compartment through the action of plasminogen activators such as t-PA and u-PA. Plasmin cleaves the fibrin network to form fibrin degradation products as illustrated in Figure 1.^32,33^ Previous studies investigating interactions between PLs, including PS, and the fibrinolytic system, have reported conflicting outcomes. ^17–21^ However, these vary substantially in methodology, including differences in the presence/absence of a fibrin surface, CaCl_2_ concentration, the incorporation of tPA directly into PL vesicles, as well the testing of non-physiological membrane compositions, containing up to 100% PS.^17–21^ To address this, our study used physiologically relevant PL compositions and found that these have no effect on the ability of plasmin to cleave its chromogenic substrate. The liposomes also had no impact on the activation of plasmin by t-PA or the dissolution of fibrin network by plasmin. Additionally, no obvious trend indicating an effect of either PS or eoxPL on the activation of plasmin by u-PA was observed. Taken together, this indicates that plasmin activation, plasmin activity, and the breakdown of fibrin by plasmin are not directly sensitive to the presence of lipid membranes composed of PC/PE, even when PS or eoxPL are included (Figure 3 A,B). In summary, isolated fibrinolytic pathways are not directly influenced by procoagulant PL, although we can’t exclude that in vivo, in the context of coagulation, changes in thrombin generation may indirectly regulate these processes.

Neither 15-HETE-PE nor 15-HETE-PC altered the activity of APC in either the absence or presence of protein S. Although APC and protein S contain Gla domains similar to pro-coagulant factors, their Kd values for binding PS/PE (10%:40%) membranes are quite different to those determined for FX, FIX and FVIIa.^40^ Specifically, while FX, FIX and FVIIa bind to the membrane with rather similar affinities, APC binds with far lower affinity (4-17 fold lower) while protein S binds far tighter (50-180 fold higher). This could provide a partial explanation for the lack of effect of eoxPL on APC/protein S, as there may be less of an impact of the wider membrane environment on the interaction of Gla domains that bind differently to the exposed PS headgroup than pro-coagulant factors.

In our study, the liposomes used were more reflective of the activated platelet surface than those previously reported and are well established to show a pro-coagulant effect, including at the concentrations and ratios tested herein.^11–13,41^ Thus, the lipid changes in the PL surface of activated platelets are unlikely to directly influence clot structure or fibrinolysis.

While our study shows no direct effect of PL on thrombin activity, clot structure, or fibrinolysis using purified systems, it is possible that these lipids could exert indirect effects through their role in supporting the coagulation cascade. Specifically, procoagulant lipids modulate thrombin generation, which in turn may influence clot structure.^26^ Differences in clot structure, and the regulation of fibrinolytic inhibitors by thrombin could result in alteration of fibrinolysis when considering the global haemostatic system.^26,32^ Platelets are well known to interact with fibrin networks and plasminogen via membrane receptors, and it is possible that this could also be influenced by aPL or eoxPL.^42,43^ Future studies could investigate whether such secondary effects of procoagulant phospholipids on clot structure and fibrinolysis occur in plasma or in vivo.

## Abbreviations

15-HETE-PC: 15-hydroxyeicosatetraenoic acid-PC
15-HETE-PE: 15-hydroxyeicosatetraenoic acid-PE
APC: activated protein C
aPL: aminophospholipids
DPPC: di-palmitoyl-phosphocholine
DSPC: 1,2-distearoyl-sn-glycero-3-phosphocholine
eoxPL: enzymatically oxidized phospholipids
FA: fatty acyl
PC: phosphatidylcholine
PE: phosphatidylethanolamine
PL: phospholipids
POPS: 1-palmitoyl-2-oleoyl-sn-glycero-3-phosphatidylserine
PS: phosphatidylserine
SAPC: 1-stearoyl-2-arachidonoyl-sn-glycero-3-phosphocholine
SAPE: 1-stearoyl-2-arachidonoyl-sn-glycero-3-phosphoethanolamine
SOPC: 1-stearoyl-2-oleoyl-sn-glycero-3-phosphocholine
SOPE: 1-stearoyl-2-oleoyl-sn-glycero-3-phosphoethanolamine
SOPS: 1-stearoyl-2-oleoyl-sn-glycero-3-phospho-L-serine
t-PA: tissue plasminogen activator
TBST: tris buffered saline with tween
u-PA: urokinase

## Acknowledgments

We would like to thank Professor. JH Morrissey (University of Michigan, USA) for interesting discussions.

## Author contributions

All authors made a significant contribution to the work reported, whether that is in the conception, study design, execution, acquisition of data, analysis and interpretation, or in all these areas; took part in drafting, revising or critically reviewing the article; gave final approval of the version to be published; have agreed on the journal to which the article has been submitted; and agree to be accountable for all aspects of the work.

## Data Availability

The datasets supporting this article can be found in the Supplementary Material. Raw absorbance data is available on reasonable request from the corresponding authors.

## References

1. Clark SR, Thomas CP, Hammond VJ, Aldrovandi M, Wilkinson GW, Hart KW, Murphy RC, Collins PW, O’Donnell VB. Characterization of platelet aminophospholipid externalization reveals fatty acids as molecular determinants that regulate coagulation. Proc Natl Acad Sci U S A. 2013;110:5875–5880. doi: 10.1073/pnas.1222419110

2. Zwaal RF, Comfurius P, Bevers EM. Lipid–protein interactions in blood coagulation. Biochim Biophys Acta. 1998;1376:433–453. doi: 10.1016/S0304-4157(98)00018-5

3. Schick PK, Kurica KB, Chacko GK. Location of phosphatidylethanolamine and phosphatidylserine in the human platelet plasma membrane. J Clin Invest. 1976;57:1221–1226. doi: 10.1172/jci108390

4. Bevers EM, Comfurius P, Zwaal RF. Changes in membrane phospholipid distribution during platelet activation. Biochim Biophys Acta. 1983;736:57–66. doi: 10.1016/0005-2736(83)90169-4

5. Antonova O, Yakushkin V, Mazurov A. Coagulation activity of membrane microparticles. Biochem Moscow Suppl Ser A. 2019;13:169–186. doi: 10.1134/S1990747819030036

6. Palta S, Saroa R, Palta A. Overview of the coagulation system. Indian J Anaesth. 2014;58:515–523. doi: 10.4103/0019-5049.144643

7. Davie EW, Ratnoff OD. Waterfall sequence for intrinsic blood clotting. Science. 1964;145:1310–1312. doi: 10.1126/science.145.3638.1310

8. Esmon CT, Suttie J, Jackson C. The functional significance of vitamin K action. Difference in phospholipid binding between normal and abnormal prothrombin. J Biol Chem. 1975;250:4095–4099. doi: 10.1016/S0021-9258(19)41391-4

9. Huang M, Rigby AC, Morelli X, Grant MA, Huang G, Furie B, Seaton B, Furie BC. Structural basis of membrane binding by Gla domains of vitamin K–dependent proteins. Nat Struct Biol. 2003;10:751–756. doi: 10.1038/nsb971

10. O’Donnell VB, Aldrovandi M, Murphy RC, Krönke G. Enzymatically oxidized phospholipids assume center stage as essential regulators of innate immunity and cell death. Sci Signal. 2019;12:eaau2293. doi: 10.1126/scisignal.aau2293

11. Thomas CP, Morgan LT, Maskrey BH, Murphy RC, Kuhn H, Hazen SL, Goodall AH, Hamali HA, Collins PW, O’Donnell VB. Phospholipid-esterified eicosanoids are generated in agonist-activated human platelets and enhance tissue factor-dependent thrombin generation. J Biol Chem. 2010;285:6891–6903. doi: 10.1074/jbc.M109.078428

12. Lauder SN, Allen-Redpath K, Slatter DA, Aldrovandi M, O’Connor A, Farewell D, Percy CL, Molhoek JE, Rannikko S, Tyrrell VJ. Networks of enzymatically oxidized membrane lipids support calcium-dependent coagulation factor binding to maintain hemostasis. Sci Signal. 2017;10:eaan2787. doi: 10.1126/scisignal.aan2787

13. Slatter DA, Percy CL, Allen-Redpath K, Gajsiewicz JM, Brooks NJ, Clayton A, Tyrrell VJ, Rosas M, Lauder SN, Watson A, et al. Enzymatically oxidized phospholipids restore thrombin generation in coagulation factor deficiencies. JCI Insight. 2018;3. doi: 10.1172/jci.insight.98459

14. Smirnov MD, Ford DA, Esmon CT, Esmon NL. The effect of membrane composition on the hemostatic balance. Biochemistry. 1999;38:3591–3598.

15. Gierula M, Salles-Crawley II, Santamaria S, Teraz-Orosz A, Crawley JT, Lane DA, Ahnström J. The roles of factor Va and protein S in formation of the activated protein C/protein S/factor Va inactivation complex. Journal of Thrombosis and Haemostasis. 2019;17:2056–2068.

16. Safa O, Hensley K, Smirnov MD, Esmon CT, Esmon NL. Lipid oxidation enhances the function of activated protein C. Journal of Biological Chemistry. 2001;276:1829–1836.

17. Exner T, Rickard K, Kronenberg H. Activation of fibrinolysis by certain phospholipids. Pathology. 1975;7:54–55. doi: 10.1016/S0031-3025(16)38930-9

18. Gerbeck C, Koppel J, Olwin JH. Effects of certain phospholipids on plasminogen activation and plasmin activity. Thromb Haemost. 1962;7:016–026. doi: 10.1055/s-0038-1655452

19. Gombás J, Tanka-Salamon A, Skopál J, Nagy Z, Machovich R, Kolev K. Modulation of fibrinolysis by the combined action of phospholipids and immunoglobulins. Blood Coagul Fibrinolysis. 2008;19:82–88. doi: 10.1097/MBC.0b013e3282f38c6f

20. Soeda S, Kakiki M, Shimeno H, Nagamatsu A. Some properties of tissue-type plasminogen activator reconstituted onto phospholipid and/or glycolipid vesicles. Biochem Biophys Res Commun. 1987;146:94–100. doi: 10.1016/0006-291X(87)90695-4

21. Varadi B, Kolev K, Tenekedjiev K, Meszaros G, Kovalszky I, Longstaff C, Machovich R. Phospholipid barrier to fibrinolysis: role for the anionic polar head charge and the gel phase crystalline structure. J Biol Chem. 2004;279:39863–39871. doi: 10.1074/jbc.M405172200

22. Bakirova D, Faizullin D, Valiullina YA, Salnikov V, Zuev YF. Effect of lipid surface composition on the formation and structure of fibrin clots. Bull Exp Biol Med. 2017;163:722–725. doi: 10.1007/s10517-017-3889-5

23. Faizullin D, Valiullina Y, Salnikov V, Zuev Y. Direct interaction of fibrinogen with lipid microparticles modulates clotting kinetics and clot structure. Nanomedicine. 2020;23:102098. doi: 10.1016/j.nano.2019.102098

24. Faizullin DA, Valiullina YA, Salnikov VV, Zuev YF. Fibrinogen Adsorption on the Lipid Surface as a Factor of Regulation of Fibrin Formation. Biophysics. 2021;66:70–76. doi: 10.1134/s0006350921010103

25. Kattula S, Byrnes JR, Wolberg AS. Fibrinogen and Fibrin in Hemostasis and Thrombosis. Arterioscler Thromb Vasc Biol. 2017;37:e13–e21. doi: 10.1161/ATVBAHA.117.308564

26. Weisel JW, Litvinov RI. Fibrin Formation, Structure and Properties. Subcell Biochem. 2017;82:405–456. doi: 10.1007/978-3-319-49674-0_13

27. Undas A, Zawilska K, Ciesla-Dul M, Lehmann-Kopydłowska A, Skubiszak A, Ciepłuch K, Tracz W. Altered fibrin clot structure/function in patients with idiopathic venous thromboembolism and in their relatives. Blood. 2009;114:4272–4278. doi: 10.1182/blood-2009-05-222380

28. Mihalko E, Brown AC. Clot Structure and Implications for Bleeding and Thrombosis. Semin Thromb Hemost. 2020;46:96–104. doi: 10.1055/s-0039-1696944

29. Carr Jr ME, Hermans J. Size and density of fibrin fibers from turbidity. Macromolecules. 1978;11:46–50. doi: 10.1021/ma60061a009

30. Ryan EA, Mockros LF, Weisel JW, Lorand L. Structural origins of fibrin clot rheology. Biophys J. 1999;77:2813–2826. doi: 10.1016/S0006-3495(99)77113-4

31. Adams TE, Huntington JA. Structural transitions during prothrombin activation: On the importance of fragment 2. Biochimie. 2016;122:235–242. doi: 10.1016/j.biochi.2015.09.013

32. Longstaff C, Kolev K. Basic mechanisms and regulation of fibrinolysis. J Thromb Haemost. 2015;13 Suppl 1:S98–105. doi: 10.1111/jth.12935

33. Risman RA, Kirby NC, Bannish BE, Hudson NE, Tutwiler V. Fibrinolysis: an illustrated review. Res Pract Thromb Haemost. 2023;7:100081. doi: 10.1016/j.rpth.2023.100081

34. Leibundgut G, Arai K, Orsoni A, Yin H, Scipione C, Miller ER, Koschinsky ML, Chapman MJ, Witztum JL, Tsimikas S. Oxidized phospholipids are present on plasminogen, affect fibrinolysis, and increase following acute myocardial infarction. J Am Coll Cardiol. 2012;59:1426–1437. doi: 10.1016/j.jacc.2011.12.033

35. Lin H, Xu L, Yu S, Hong W, Huang M, Xu P. Therapeutics targeting the fibrinolytic system. Exp Mol Med. 2020;52:367–379. doi: 10.1038/s12276-020-0397-x

36. Zabczyk M, Ariens RAS, Undas A. Fibrin clot properties in cardiovascular disease: from basic mechanisms to clinical practice. Cardiovasc Res. 2023;119:94–111. doi: 10.1093/cvr/cvad017

37. Morgan AH, Hammond VJ, Morgan L, Thomas CP, Tallman KA, Garcia-Diaz YR, McGuigan C, Serpi M, Porter NA, Murphy RC, et al. Quantitative assays for esterified oxylipins generated by immune cells. Nat Protoc. 2010;5:1919–1931. doi: 10.1038/nprot.2010.162

38. Smirnov MD, Esmon CT. Phosphatidylethanolamine incorporation into vesicles selectively enhances factor Va inactivation by activated protein C. Journal of Biological Chemistry. 1994;269:816–819.

39. Maskrey BH, Bermudez-Fajardo A, Morgan AH, Stewart-Jones E, Dioszeghy V, Taylor GW, Baker PR, Coles B, Coffey MJ, Kuhn H. Activated platelets and monocytes generate four hydroxyphosphatidylethanolamines via lipoxygenase. J Biol Chem. 2007;282:20151–20163. doi: 10.1074/jbc.M611776200

40. Medfisch SM, Muehl EM, Morrissey JH, Bailey RC. Phosphatidylethanolamine-phosphatidylserine binding synergy of seven coagulation factors revealed using Nanodisc arrays on silicon photonic sensors. Sci Rep. 2020;10:17407. doi: 10.1038/s41598-020-73647-3

41. Billy D, Willems GM, Hemker HC, Lindhout T. Prothrombin contributes to the assembly of the factor Va-factor Xa complex at phosphatidylserine-containing phospholipid membranes. J Biol Chem. 1995;270:26883–26889. doi: 10.1074/jbc.270.45.26883

42. Buitrago L, Lefkowitz S, Bentur O, Padovan J, Coller B. Platelet binding to polymerizing fibrin is avidity driven and requires activated αIIbβ3 but not fibrin cross-linking. Blood Adv. 2021;5:3986–4002. doi: 10.1182/bloodadvances.2021005142

43. Whyte C, Morrow G, Baik N, Booth N, Jalal M, Parmer R, Miles L, Mutch N. Exposure of plasminogen and a novel plasminogen receptor, Plg-RKT, on activated human and murine platelets. Blood. 2021;137:248–257. doi: 10.1182/blood.2020007263

